# Male attraction to short-range multimodal gyne signals in bumble bees

**DOI:** 10.1101/2025.06.10.658960

**Authors:** Sarah K. Spence, Etya Amsalem

## Abstract

Mating is a crucial event in animal life, with significant implications for individual fitness. Insects have evolved complex mating systems and behaviors, such as pheromone signaling and elaborate courtship rituals, that convey vital information about mates’ species, quality, and receptiveness, to optimize reproductive success. These signals are especially critical in social species where a single female found a colony following a single event of mating.

In this study, we investigate the short-range signals produced by gynes (newly emerged queens) and their role in mating in *Bombus impatiens*. Mating in bumble bees is facilitated by males marking mating sites with long-range labial gland pheromones, which attract gynes. It is assumed that gynes produce short-range signals necessary to initiate mating. While male bumble bee sex pheromones have been extensively studied across many species, the short-range signals produced by gynes remain poorly understood. Our findings show that males rely on multimodal signals, composed of visual, vibrational, and context-dependent sex pheromones produced by gynes. Choice bioassays designed to isolate the different signals produced by gynes indicate that visual cues are essential but not sufficient for successful mating, and compounds from the labial and dufour glands likely play a role in attracting males. Our results also highlight the importance of vibrational signals for male attraction. Together, these findings underscore the role of gynes in regulating mating and the significance of context in eliciting male responses and mate selection through various signals.

## INTRODUCTION

Mating is a pivotal event in the life cycles of many organisms, influencing reproductive success and genetic diversity for both parents and offspring. In social insects, mating holds particular significance as it shapes colony structure, genetic diversity, and reproductive strategies. For instance, monandrous queens (that mate only once) produce colonies with high relatedness among workers, which reduces reproductive conflict and enhances cooperation [1]. Additionally, female preference for male traits, such as distinct abdominal spot patterns in *Polistes dominula*, can shape colony genetics and social dynamics [2], while the number of matings per female may impact the colony’s sex ratio [3]. For social insect females, mating is often a single event, where they store sperm in a specialized pouch to fertilize eggs throughout their lives before establishing a colony, making that one mating event disproportionately impactful for the lives of many individuals. Furthermore, in species where monandrous females and polygynous males coexist, a common mating system in social insects [4], along with a male-biased sex ratio, male competition and female selectivity intensify, strengthening sexual selection pressures, mating costs and reliance on mating signals [5]. To optimize these fitness consequences, social insects have evolved diverse signals to facilitate mate recognition and selection.

Choosing a mate is a complex process that involves multiple challenges, such as selecting the correct species and sex, determining receptiveness to mating, and evaluating mate quality. In social insects, this complexity is heightened by signals that also convey information about sociality, relatedness, and caste [6, 7]. To facilitate mating, insects employ diverse visual [8], chemical [9, 10], and auditory signals [11]. Examples include the courtship dances of male fruit flies [12], vibrational "drumming" in stoneflies [13], and the mating calls of male crickets and cicadas [14, 15]. Furthermore, insects have extensively adapted the use of pheromones, with sex pheromones identified in numerous species, including moths, beetles, cockroaches, wasps, and thrips [16–20].

Many species utilize multiple signals simultaneously during mating, with one signal providing context to enhance the accuracy of another. For instance, male Osmia bees combine chemical and vibrational signals by vibrating their thorax while rubbing their antennae and bodies on the female’s antennae [21]. Similarly, male parasitoid wasps (*Leptopilina heterotoma*) are only attracted to female sex pheromones when combined with wing-fanning behavior [22]. In honeybees, the queen emits sex pheromones from her mandibular glands to signal receptiveness during the nuptial flight [23, 24], and the same compound (9-ODA) later functions to induce retinue behavior in worker bees after colony establishment [25]. Likewise, in *Rhytidoponera metallica* ants, females release sex pheromones while lowering the head and thorax and raising the gaster at the nest entrance [26]. The use of multimodal signals extends beyond mating to other critical behaviors. Bumble bees learn to identify flowers more effectively when both olfactory and visual cues are presented together rather than individually [27]. Aphaenogaster ants elicit cooperative prey transport by combining poison gland secretions with stridulation [28] and carpenter bees struggle to locate their nests when the visual or olfactory markers of the site are altered [29]. The diversity of signals and their context-dependent roles complicates their identification but provides valuable insights into the evolutionary pressures shaping complex mating systems and behaviors across species.

In this study, we investigated the short-range signals used by bumble bee gynes (*Bombus impatiens*) to attract mates, with a specific focus on the multimodal signaling strategies of males. Bumble bees are annual, social insects that produce reproductive individuals (sexuals) late in the summer [30, 31]. Male bumble bees mate multiple times and die shortly afterward, while gynes mate only once and then enter winter diapause. Mating in bumble bees involves long-range signals produced by males, likely combined with short-range signals from gynes. Male bumble bees establish and defend territories to attract gynes, marking these areas with labial gland secretions deposited on leaves and stones [32–36]. These signals are paired with species-specific behaviors, such as patrolling (marking multiple sites and flying circuits around the territory), perching (marking a single site and waiting nearby) or waiting at the nest entrance of a colony for gynes to leave. These strategies rely heavily on visual cues, supported by males’ relatively large eyes [34]. While male signaling and mating behaviors have been extensively studied, the signaling role of gynes remains poorly understood, highlighting a critical gap in our knowledge of bumble bee mating dynamics.

Previous studies on bumble bee mating have focused on the sex pheromones produced by gynes, identifying potential ester compounds in their dufour and labial glands that correlate with sexual maturity in both *Bombus terrestris* [37, 38] and *Bombus impatiens* [39, 40]. Additionally, extracts from the cuticular surface and the entire head of *Bombus terrestris* gynes have been shown to elicit mating behaviors such as touching and mounting, with the mandibular glands assumed to be the source of these signals [41, 42]. Despite these findings, no specific compounds have been conclusively identified, and the signals governing mating remain largely unknown. To address these gaps, we conducted a comprehensive investigation into the signals produced by bumble bee gynes to attract males. This included testing male responses to visual, chemical (from various glands), and vibrational signals produced by gynes. Through a series of olfactometer bioassays, we isolated and evaluated the individual and combined effects of these signals.

## METHODS

### Bees

Colonies of bumble bees (*Bombus impatiens*) were obtained from Koppert Biological Systems and kept in environmental chambers at 28-30°C and 60% relative humidity and served as a source for males, gynes and workers. The colonies were kept in darkness, were handled under red light and supplied with *ad libitum* 60% sucrose solution and fresh pollen collected by honeybees (also purchased from Koppert). Males were collected upon emergence from colonies or from small queen-less cages (6 x 6 x 10 cm) containing three workers and were kept in groups of 1-5 until the age of 6-8 days. Newly emerged gynes were reared in cages containing 1-3 related gynes until the age of 6-11 days. These age ranges were found optimal for mating in a previous study from our lab [43]. Workers were sampled directly from the colonies. All cages were reared under the same conditions as the colonies.

### Overview of the experimental design

We conducted 5 series of bioassays in which males were placed in a two-choice olfactometer (see Figure 1) containing two choices. In the first set, used to optimize the experimental set up (227 bioassays), we offered males the choice between two empty cups (examining their preference to left/right), an empty cup vs. pollen (examining their preference to a positive stimulus (food) to assess the overall response rate in our system); and between the two solvents used in the study on a glass coverslip (to assess whether any of them is preferred by bees). In the second set (147 bioassays), we offered the males the choice between live gynes and live workers under either red or ambient light, and between dummy gynes and workers (see below for “dummy preparation”). Complementary to the second set, we examined the physical interaction of males with freshly frozen and dummy bees from the second set (59 bioassays). In the third set (166 bioassays), we offered the males the option between a solvent control and a chemical extract from the gyne’s cuticular lipids, dufour, labial and mandibular glands (see below for “extract preparation”). All extracts were applied to a glass coverslip. In the fourth set (172 bioassays), we repeated the bioassays from the third set, but this time we applied the extracts to dummy gynes. In the fifth and final set (170 bioassays), we again repeated the third set, but this time we applied the extracts on top of a newly emerged worker. We collected data on the choice made by males, the number of interactions they made with bees and the overall response rate in all bioassays. Overall, we conducted 941 choice bioassays where both males and choices used only once (excluding the 59 bioassays that are complementary to the second set).

**Figure 1.**
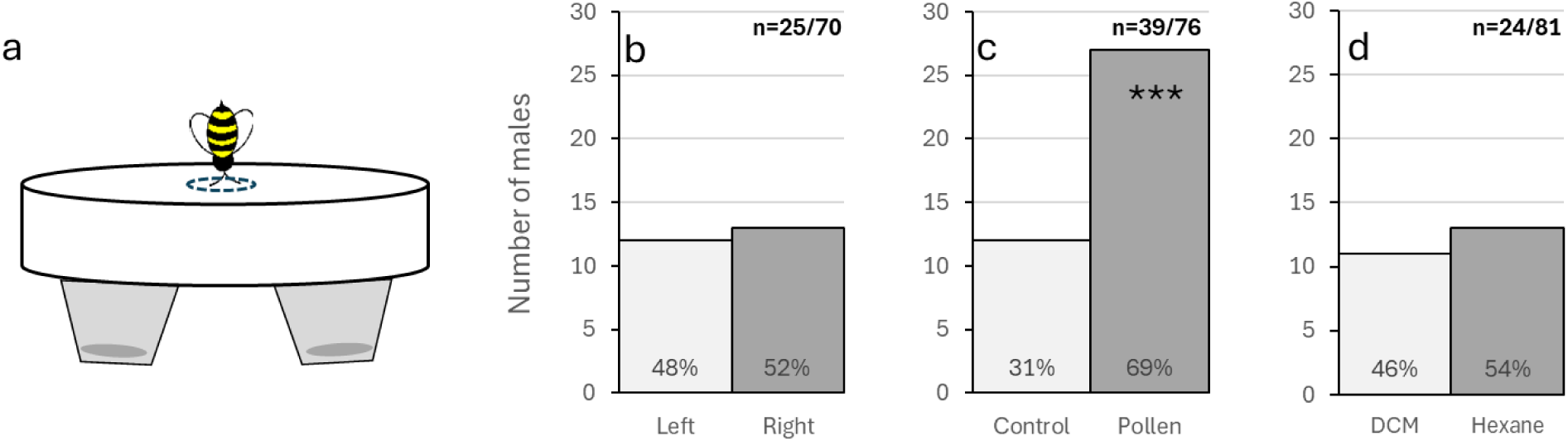
A validation of the experimental set up. Across all experiments, males were placed in a two-choice olfactometer (a) and provided with a choice composed of dead or live bees and various chemical extracts in a solvent. To set the baseline for male response in these experiments, we examined males’ preference to a direction (left vs. right) (b), a positive stimulus (pollen vs. control) (c), and the two solvents we used in the study (Dichloromethane (DCM) vs. hexane). Males did not have a preference to a specific direction or a solvent but were two times more likely to choose the positive stimulus (pollen) over the control. The sample size (n=x/y) is presented as the number of responding males/overall sample. The figures include only responding males. The response rate was higher (51%) when a positive stimuli was offered vs. options which the male had no preference to (31-36%). Asterixis denote significant differences at *p* < 0.05 (*), *p* < 0.01 (**), *p* < 0.001 (***).

### Extract preparation

Cuticular lipid extracts were obtained from 6-11 days old gynes (86 extracts in total). To collect the extract, a live gyne was placed in a 20 mL scintillation vial for 10 minutes as in [44] and the vial was rinsed with 2 mL dichloromethane. Gynes were returned to their cage and additional extracts were collected in the following days until gynes reached the age of 11 days. The vials were evaporated, and every 10-25 extracts of gynes were combined and redissolved in dichloromethane to achieve a concentration of 10 µl per one queen equivalent extract.

Dufour, labial and mandibular glands extracts were obtained from approximately 150 gynes at the age of 6-11 days. Gynes were frozen at -80°C and dissected as follows. The abdomen was sliced opened under a stereomicroscope and the ovaries were removed and measured (*i.e.*, to ensure gynes didn’t activate their ovaries, see below). The dufour’s gland was then extracted from the abdomen as in [38] and the labial and the mandibular glands were extracted from the head as in [39, 45]. Dissected glands were removed at the base of the gland to avoid puncture. Glands were placed on an ethanol-cleaned pin and transferred immediately into a glass vial containing 200 µl of dichloromethane (DCM). Each vial contained 8-20 same type glands and was marked for the number of glands in it. The vials were stored at -20°C until used. Extracts were nearly evaporated and redissolved in DCM or hexane solvent in a solution of 10 µl/gland. DCM was used in all experiments excluding the extracts applied on top of live workers due to its toxicity to bees. Extracts were used in all bioassays in a dose equivalent to one gyne and a volume of 10 µl. The extracts or the solvent were applied to either a glass coverslip, a dummy gyne, or a newly emerged worker. The solvent was let to evaporate for 5-10 minutes before the male was introduced into the olfactometer. Newly emerged workers were sampled from their colony 10-15 prior to the onset of the bioassay and the volume applied to their thorax was reduced to one gyne equivalent in 2 µl. The extracts were slowly applied to the workers in 0.5 µL increments. Every callow (newly-emerged worker) was used only once.

To ensure that all the extracts came from virgin gynes that did not activate their ovaries, we further measured the average terminal oocyte of gynes during dissection as in [38]. Normally, all gynes at that age range have inactivated ovaries, but under captivity some gynes will activate their ovaries. If the average terminal oocyte of a gyne was greater than 2 mm (which indicates activated ovaries), the gyne was discarded and glands were not dissected out.

### Preparation of dummies

Dummy bees were made using frozen queens and workers that were kept at -80° C until processing. To remove all scents, bees were placed in a Soxhlet extraction column with 2:1 chloroform to methanol solution for at least 4 hours. A small cut was made horizontal to each bee’s stinger before extraction. The dummies were rinsed with 70% ethanol solution and distilled water, air dried in a fume hood, and blow-dried to prevent a matted appearance. Bees were placed in the oven at 65° C for about 5-7 days and stored at room temperature in plastic tubes until use. When used in bioassays, dummies were made to stand as realistically as possible. The extracts or the solvent were applied to the thorax and the abdomen of the dummy, and the solvent was allowed to evaporate before introducing the male. Every dummy was used only once.

### Olfactometer Bioassays

All bioassays took place in a two-choice olfactometer (Figure 1a) conducted between 4 to 8 pm. Excluding two bioassays that were conducted in the environmental chamber under red light, all other bioassays were conducted at RT under ambient light. Males were introduced into the olfactometer and were given 30 minutes to make a choice. The olfactometer is built in a way that once a male makes a choice and enters one of the cups, he can no longer go back.

### Contact behavior of males

In several bioassays (Figure 2d), we also tested the contact behavior of males with the choices. After olfactometer bioassays were completed, new males were placed in a large plastic arena to acclimate for 15 minutes. The live gynes and workers from the second set of bioassays were frozen on dry ice and reused by placing each choice on one side of the arena. The male was given 50 min to interact with the bees, during which his behavior was video-recorded and the percentage of time he interacted with each of the choices (sum of touching, climbing and mounting incidents) was calculated. Touching was defined as any contact, climbing was defined as more than two legs on the object or being over the object, and mounting was defined as pulling or pushing movements towards the bee. We repeated this experiment also with the worker and gynes dummies from the second set of bioassays.

**Figure 2.**
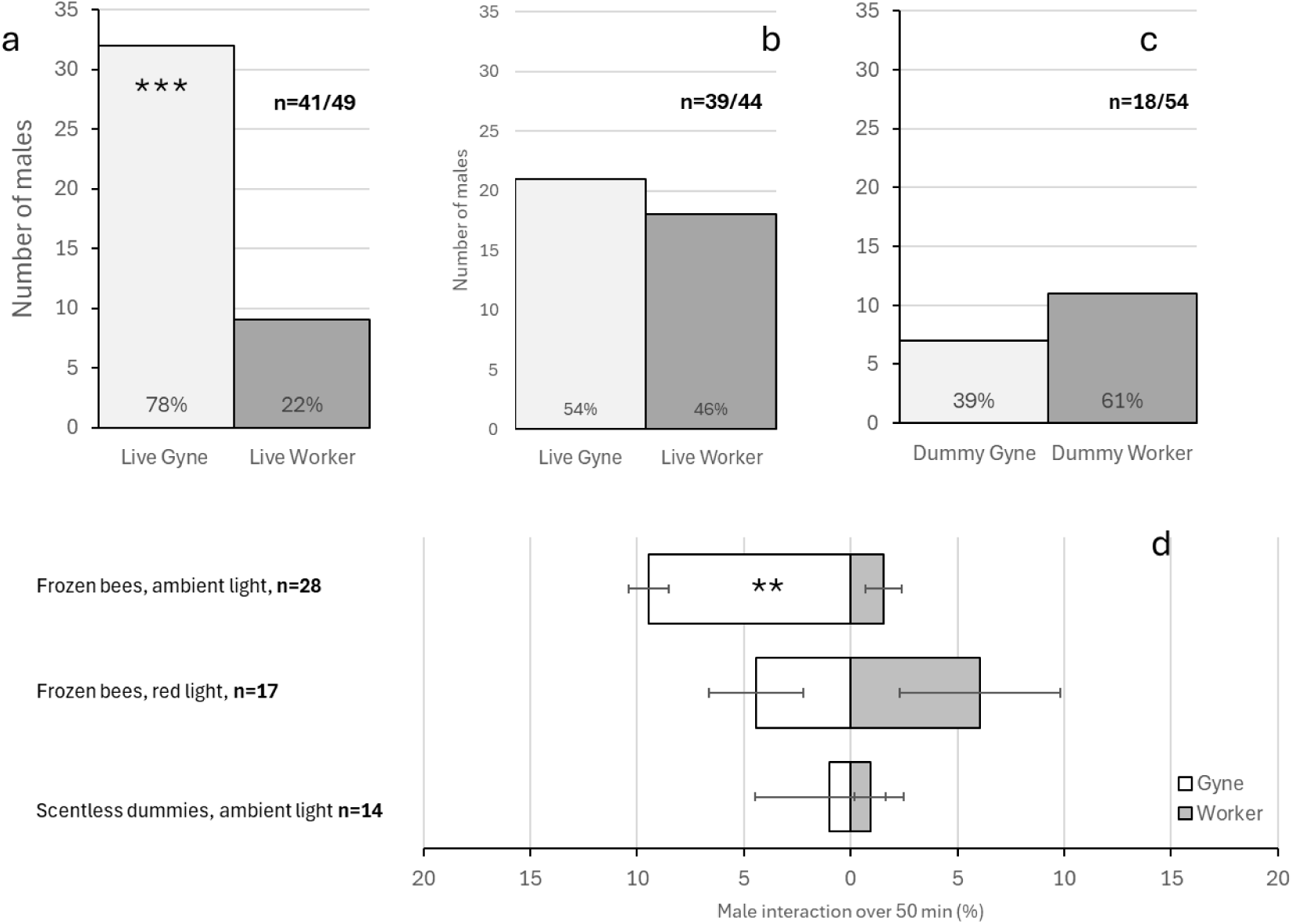
Short range attraction of males in response to visual signals of bees. Males were offered the choice between live worker vs. live gyne under either ambient light (a) or red light (b) and favored gynes only under ambient light. Male response to visual signals alone was tested under ambient light using scentless dummies of workers and gynes (c) to which males showed no preference. The sample size (n=x/y) is presented as the number of responding males/overall sample. Figures a-c include only responding males. Freshly frozen samples from a-b and dummies were placed in an arena containing a single male for 50 minutes. The physical interactions of males with dead bees matched their performance in the olfactometer with overall more interactions with dead gynes in ambient light (d). Asterixis denote significant differences at p < 0.05 (*), p < 0.01 (**), p < 0.001 (***).

### Statistical analysis

Data analysis and visualization were conducted using R studio and JMP pro-18. Bioassays were compared in R using the Chi-Square Test of Independence that tests whether the observed frequency of choices in each arm differs from what would be expected if there were no preference. The count data of male contact with frozen/dummy females were compared using GLM with Poisson distribution and log as link function with overdispersion and followed by contrast comparisons. The treatment, female type and their interaction were used as fixed variables, and the interaction time of males was used as the dependent variable. The correlation between the p-value and the response rate was tested in JMP using Pearson correlation. Data are presented as mean ± SE. Significant differences were accepted at α=0.05.

### Ethical note

A license or certificate was not required for experiments with bees. However, all experiments adhered to international ethical standards for animal care and use. To minimize stress, bees were handled under red light during collection, housing, and testing. They were housed in appropriately sized cages for their group size and provided with unlimited food throughout the experiment. Euthanasia was carried out by placing bees in a -80°C freezer.

## RESULTS

Across all experiments, males were tested for their attraction to various signals in a two-choice olfactometer (Figure 1a). To validate the experimental setup, we tested male attraction to a direction (left vs. right), a positive stimulus (pollen vs. empty cup) and to the solvents we used in the study (dichloromethane vs. hexane). Males showed no preference between left and right (ꭓ^2^ = 0.04, *df* = 1, p= 0.84, n=25/70 (the n is described using two parameters: the number of responders followed by the total number of bioassays), response rate 36%, Figure 1b). Males were attracted to the pollen vs. the empty cup (ꭓ^2^= 5.77, *df* = 1, p= 0.02, n=39/81, response rate 51%, Figure 1c) and had no preference between the solvents (ꭓ^2^= 0.166, *df* = 1, p= 0.68, n=25/81, response rate 30%, Figure 1d). The response rate aligned with the interest males had in the options offered to them with 30-36% response rate when they had no preference and 51% response rate when they did.

When males were offered the choice between a live worker and a live gyne, they showed a preference towards gynes under ambient light (ꭓ^2^ = 12.9, *df* = 1, p= 0.0003, n=41/49, response rate 84%, Figure 2a) and a lack of preference under red light (ꭓ^2^ = 0.23, *df* = 1, p= 0.63, n=39/44, response rate 87%, Figure 2b). When we offered males with a dummy worker vs. a dummy gyne under ambient light in order to eliminate the auditory/vibration and chemical signals coming from live bees, this has resulted in males showing no preference with overall low response rate (ꭓ^2^ = 0.89, *df* = 1, p= 0.35, n=18/54, response rate 33%, Figure 2c). Freshly frozen samples from 1a-b and dummies were introduced to males in a follow up experiment where they were placed in an arena and the physical interactions of the males with them were video-recorded for 50 minutes. The performance of males in the arena with freshly dead bees matched the performance in the olfactometer with more interactions of males with dead gynes over workers under ambient light (GLM, ꭓ^2^= 14.2, *df* = 5, p= 0.01, n=118 followed by contrast comparison between gyne and worker under ambient light: ꭓ^2^= 8.08, *df* = 1, p= 0.004, n=56).

Next, we isolated the putative chemical signal by offering males with a choice between a solvent control and an extract of gynes’ cuticular lipids, dufour, labial, and mandibular glands. The extract was placed on a glass coverslip and the bioassays were conducted under ambient light. Males showed no preference between the solvent and the cuticular secretion (ꭓ^2^= 0, *df* = 1, p= 1, n=12/40, response rate 30%, Figure 3a), showed a slight but insignificant preference to the dufour’s gland over the solvent (ꭓ^2^= 3.2, *df* = 1, p= 0.07, n=20/51, response rate 39%, Figure 3b), had a preference to the solvent over the labial gland (ꭓ^2^= 3.76, *df* = 1, p= 0.05, n=13/46, response rate 28%, Figure 3c), and showed no preference between the solvent and the mandibular gland (ꭓ^2^= 2, *df* = 1, p= 0.16, n=8/29, response rate 28%, Figure 3d).

**Figure 3.**
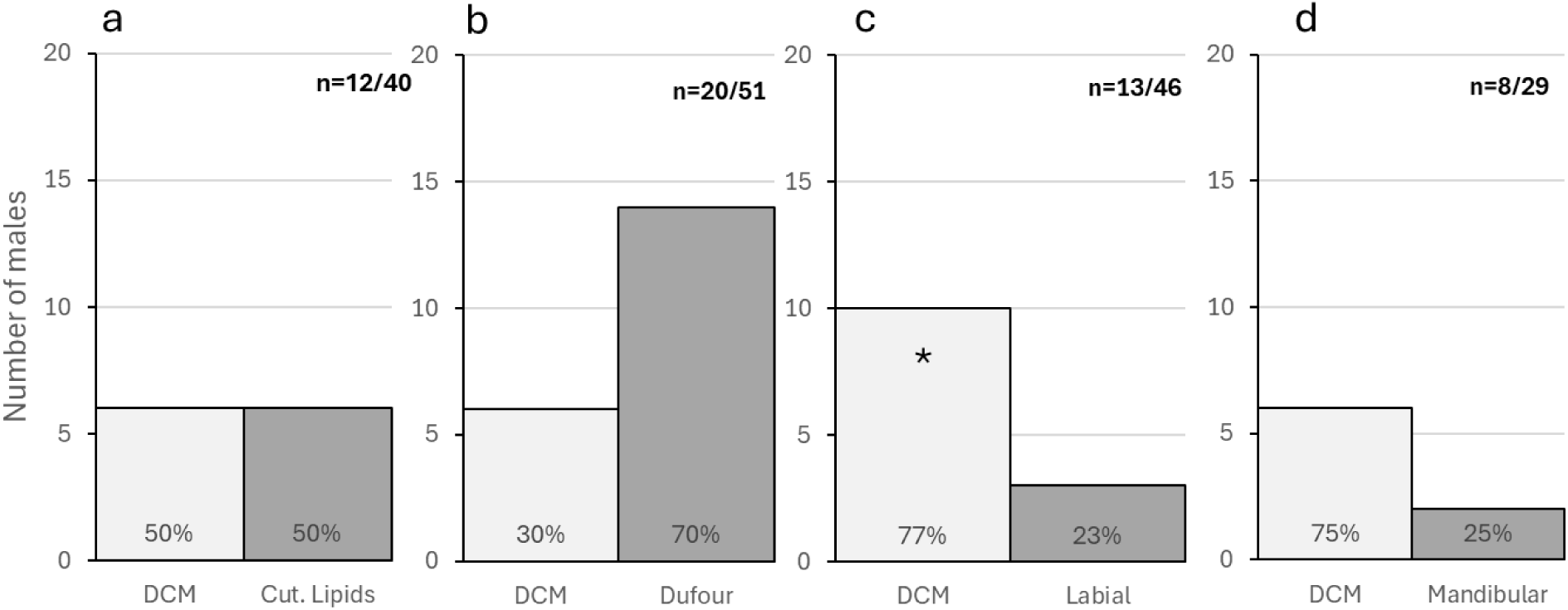
Short term attraction of males in response to gynes’ extracts applied on coverslips. Males were offered the choice between dichloromethane solvent control (DCM) vs. one of the following gynes’ extracts: cuticular lipids (a), dufour (b), labial (c) and mandibular (d) glands. Males showed a slight, insignificant preference to dufour’s gland over the solvent, and a significant preference to solvent over the labial glands. No preference was found toward the cuticular lipids and mandibular gland extracts. The sample size (n=x/y) is presented as the number of responding males/overall sample. The figures include only responding males. Asterixis denote significant differences at *p* < 0.05 (*), *p* < 0.01 (**), *p* < 0.001 (***).

In the next series of bioassays, we combined the putative chemical signal with a visual signal by offering males a choice between extracts vs. solvent on dummies. Here too we included extracts of the cuticular lipids, dufour, labial and mandibular glands and conducted assays under ambient light. Males showed a preference to the labial glands on dummies over the solvent (ꭓ^2^= 5, *df* = 1, p= 0.02, n=20/46, response rate 44%, Figure 4c), and did not show a preference towards the cuticular lipids (ꭓ^2^= 1.33, *df* = 1, p=0.25, n=12/46, response rate 26%, Figure 4a), dufour (ꭓ^2^= 2.25, *df* = 1, p=0.13, n=16/51, response rate 31%, Figure 4b) and mandibular glands (ꭓ^2^= 1.28, *df* = 1, p=0.25, n=7/29, response rate 24%, Figure 4d).

**Figure 4.**
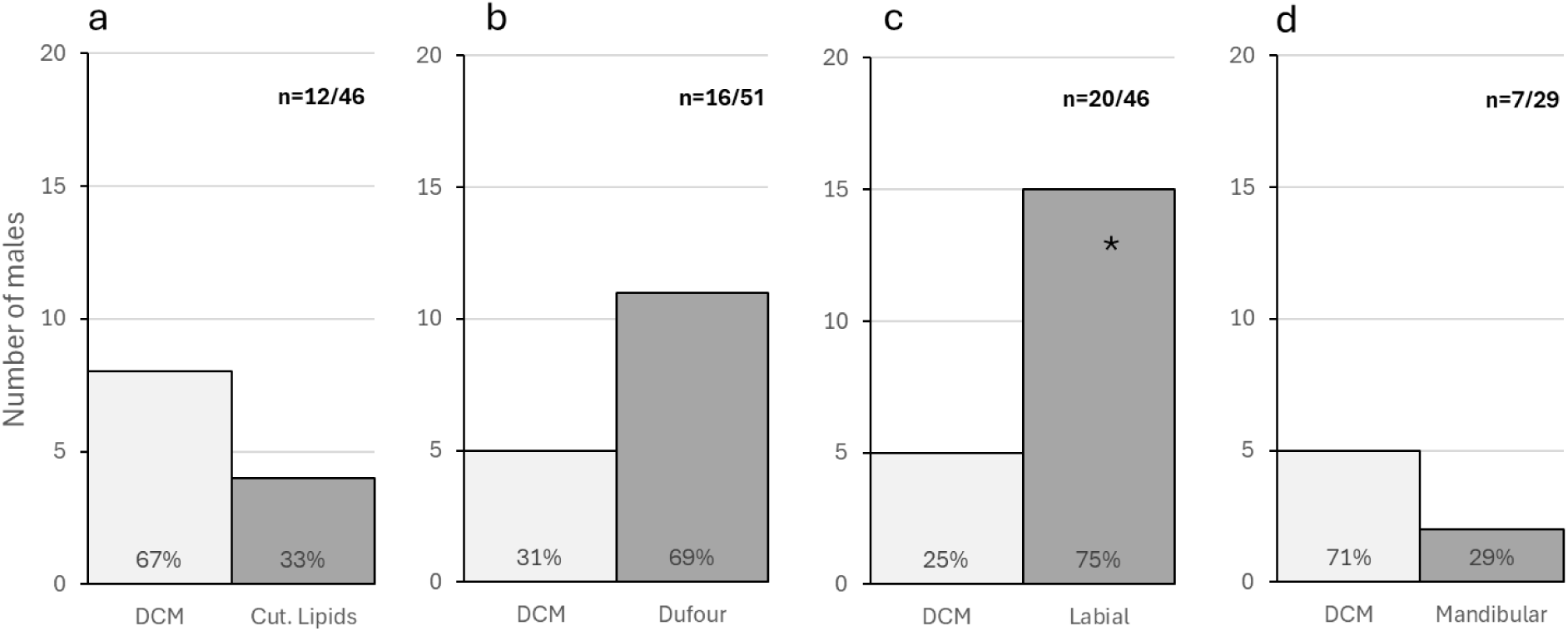
Short term attraction of males in response to gynes’ extracts applied to dummies. Males were offered the choice between a solvent control (dichloromethane) vs. one of the following gynes’ extracts applied to dummy gynes: cuticular lipids (a), dufour (b), labial (c) and mandibular (d) glands. Males showed a preference to dufour and labial glands over the solvent, and no preference in all other cases. The sample size (n=x/y) is presented as the number of responding males/overall sample. The figures include only responding males. Asterixis denote significant differences at *p* < 0.05 (*), *p* < 0.01 (**), *p* < 0.001 (***).

In the final set of bioassays, we combined the putative chemical signals with a partial visual and vibrational signal by applying the extracts on a live newly emerged worker. To isolate the vibrational signal, we first offered males the choice between a live and a dummy worker and found a significant preference to a live worker (ꭓ^2^ = 12.8, *df* = 1, p = 0.0003, n=20/42, response rate 48%, Figure 5a). When we applied the gynes’ extracts to the newly-emerged workers, we observed a significant preference to the solvent over the labial glands (ꭓ^2^= 5.26, *df* = 1, p= 0.02, n=23/40, response rate 58%, Figure 5c) and no preference by males in the bioassays with dufour (ꭓ^2^ = 1.5, *df* = 1, p = 0.22, n=24/52, response rate 46%, Figure 5b) or the mandibular glands (ꭓ^2^ = 0.53, *df* = 1, p = 0.46, n=17/36, response rate 47%, Figure 5d).

**Figure 5.**
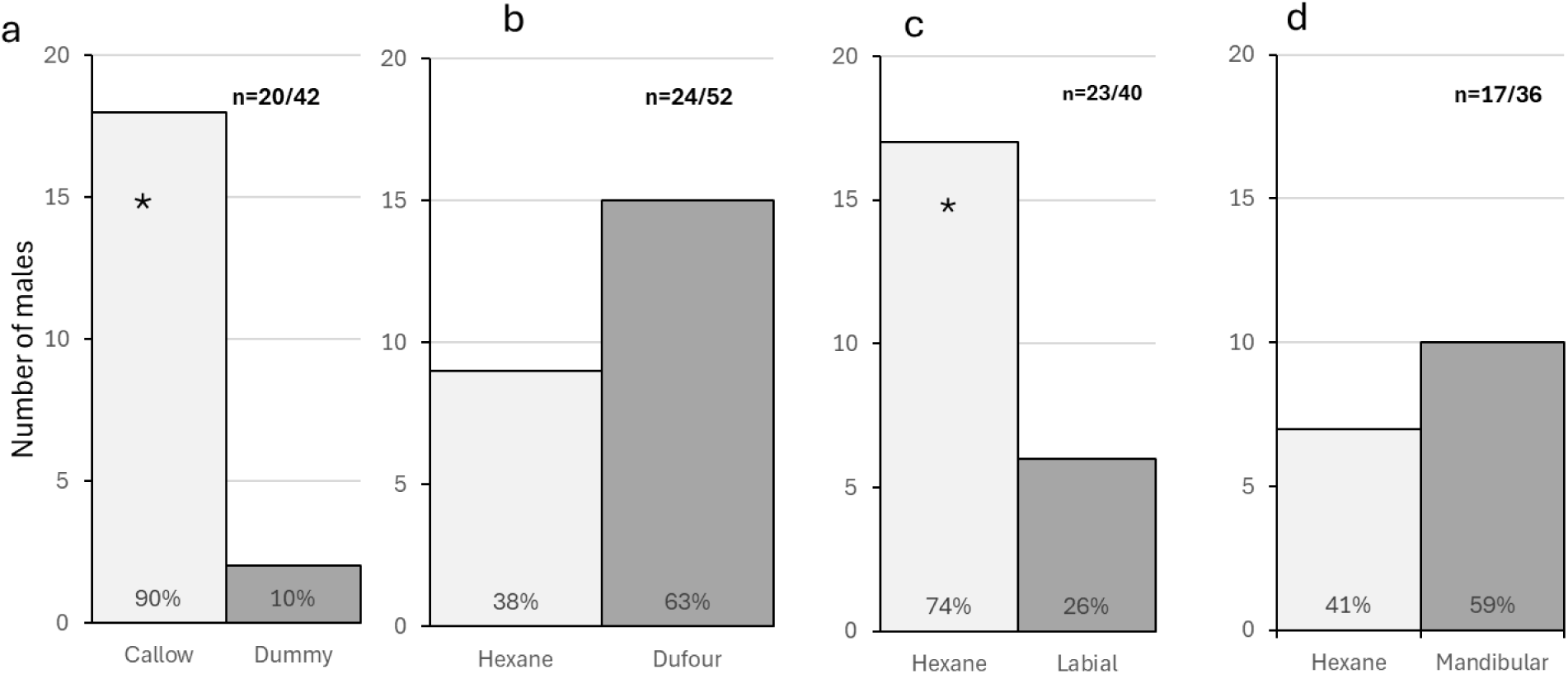
Short term attraction of males in response to gynes’ extracts applied to newly emerged workers. males were more attracted to live newly emerged worker vs. a dummy worker (a), suggesting the involvement of a vibrational signal. Males were therefore offered the choice between a solvent control (Hexane) vs. one of the following gynes’ extracts on top of a live newly emerged worker: dufour (b), labial (c) and mandibular gland (d). Males showed a preference to the solvent over the labial gland. The sample size (n=x/y) is presented as the number of responding males/overall sample. The figures include only responding males. Asterixis denote significant differences at p < 0.05 (*), p < 0.01 (**), p < 0.001 (***).

While male choice was the best way to assess male preference, we found that the response rate was also indicative of male preference, with overall higher response rate when males had a preference. To test that, we plotted the p-value of all the bioassays that were conducted under ambient light against the response rate. When we used all data (n=17), the correlation was high (r=-0.39) but insignificant (p=0.11). When we used only the bioassays with significant p-values (n=6), the correlation was very strong and nearly significant (r=-0.76, p=0.07).

## DISCUSSION

In this study, we examined male attraction to visual, chemical, and vibrational signals produced by gynes. Our findings provide key insights into the short-range signaling mechanisms regulating mating in *Bombus impatiens*. First, both male choice and overall response rate offer valuable indicators of male behavior, as a lack of preference often correlated with a low response rate. Second, male short-range attraction depended on multiple signals working together in a context-dependent manner. Visual cues were the most crucial, followed by chemical signals likely produced in the dufour and/or labial glands, with vibrational signals also playing a possible role.

The use of multimodal signals to facilitate mating may help ensure its effectiveness, especially in social insects where gynes mate only once in their lifetime and produce hundreds of offspring. The combination of signals may serve as a contextual layer, increasing the likelihood of signal accuracy. Multiple factors may contribute to this reliance on multimodal signaling in bumble bees. First, the power dynamics between males and gynes play a role; gynes are three times larger than males and equipped with a sting. In laboratory settings, we frequently observe gynes displaying aggression toward males attempting to mate, coupled with an overall low receptivity of gynes to mating. This reliance on multiple signals may thus serve as a form of defense for males. Additionally, bumble bee populations often have a male-biased sex ratio [46], leading to intense competition among males, many of whom may fail to mate. Refining mating success by responding to multiple signals could increase their chances of successful mating. Lastly, although males, who can mate multiple times, are expected to be less selective than single-mated females [47], males might still assess gyne quality before mating. Visual, vibrational, and chemical signals could serve as honest indicators of a gyne’s ability to successfully overwinter and initiate a nest. In previous study [39], we show for instance that the labial gland secretion of *Bombus impatiens* gynes is indicative of their age and reproductive status.

Bumble bee males are known for their enhanced vision and large eyes, which is even more enhanced in species where males are exhibiting perching behavior [48]. Indeed, males showed no attraction to gynes when visibility was limited (e.g., under red light), indicating the importance of visual cues. However, visual signals alone were insufficient, as males showed no preference for dummies, suggesting that multiple signals are necessary for mating facilitation. When we isolated chemical compounds (gland extracts), we found evidence that both the labial and dufour glands likely play a role in mating mediation. Bioassays revealed that males displayed a slight, insignificant attraction to the dufour’s extract and a significant aversion to the labial extract when presented on cover slips. Interestingly, when these extracts were applied to dummy gynes, males were attracted to the labial extract but showed no preference for the dufour’s extract. When the extracts were applied to callow workers, males again showed a preference for the labial gland. Previous studies from our lab (Derstine, Villar et al., 2021; Orlova, Villar et al., 2022) identified gyne-specific ester compounds in gland compositions that may act as sex pheromones, though the chemical structures remain partially characterized, and their functionality has not yet been tested. Although gyne chemical signals induced a relatively weak and context-dependent response, they were insufficient to replicate the full effect of a live gyne, suggesting that while chemical compounds are important, they alone do not suffice.

In the related species *Bombus terrestris*, the gyne sex pheromone is thought to originate from the mandibular gland [42], based on bioassays with the whole head, the cuticular extract and the similarity between head and mandibular gland extracts. However, in *Bombus impatiens*, the mandibular gland does not appear to elicit a similar response. While this may indicate species differences, it is also possible that in *Bombus terrestris*, the mating signal originates from the labial glands, which are also located in the head.

A separate test of vibrational signals highlighted their significance, as males preferred live gynes over workers and live workers over dummies, suggesting they respond to vibrations in a quantitative and relative manner. Gynes likely produce a stronger buzzing sound than workers, which is attractive to males. However, in its absence, males are still drawn to the lower buzzing sound likely produced by a worker. In captivity, males show a preference for mating with gynes but will attempt to mate with workers as well [49].

When all signals (visual, chemical extracts, and vibrational) were combined, male responses remained relatively weak, showing minor, non-significant attraction to the dufour’s extract and avoidance of the labial extract. This could be due to several factors. First, the use of callow workers instead of gynes in the final set of bioassays may have provided only partial visual and vibrational signals. Second, the dose of labial and dufour gland extracts may have been insufficient to elicit a response, especially given the suspected low volatility of the long-chain ester compounds considered as sex pheromones, combined with the lack of direct male-extract contact. Future studies should focus on specific compounds from these glands and pair them with more precise visual and vibrational signals to determine the exact modalities males use for mating. It would also be useful to combine the secretion of the glands likely involved in attraction and see their effect on male attraction and mating in concert with other stimuli.

### Conclusions

Our data indicate that male bumble bees use multimodal signals to facilitate short-range mating interactions. Visual, chemical, and vibrational signals all play critical roles, though none are sufficient independently. Compounds from the labial and dufour glands of gynes likely contribute to mating mediation. Multimodal signaling can enhance signal reliability, especially in high-stakes mating events typical of social insects

## STATEMENTS AND DECLARATIONS

### Availability of data and material

The data generated in the current study are available as a supplementary material.

### Author contribution

SS conducted the experiments, analyzed the data and wrote the first draft. EA designed the study and wrote the paper together with SS. All authors gave final approval for publication and agree to be held accountable for the work performed therein.

